# Overexpression of Channelrhodopsin 2 interferes with the GABAb receptor-mediated depression of GABA release from the somatostatin-containing interneurons of the prefrontal cortex

**DOI:** 10.1101/168245

**Authors:** Lei Liu, Wataru Ito, Alexei Morozov

## Abstract

Region and cell-type restricted expression of light activated ion channels is the indispensable tool to study properties of synapses in specific circuits and to monitor synaptic alterations by various stimuli including neuromodulators and behaviors, both ex vivo and in vivo. These analyses require the light-activated proteins or viral vectors for their delivery that do not interfere with the phenomenon under study. Here, we report a case of such interference in which the high-level expression of Channelrhodopsin-2 (ChR2) introduced in the somatostatin-positive GABAergic neurons (SOM-INs) of the dorsomedial prefrontal cortex (dmPFC) by an adeno-associated virus vector (AAV) weakens the presynaptic GABAb receptor-mediated suppression of GABA release.

## Introduction

In circuit analyses, channelrhodopsin-2 (ChR2) and its’ analogue opsins have been used to identify long-range synaptic connections (1) and to compare strength of synapses among distinct classes of neurons within a local microcircuit (2). Moreover, the opsins enabled investigation of plasticity in specific synapses in response to various factors including neuromodulators (3), behavioral experience (4–6) and genetic mutations (7, 8). Given the evidence that presynaptically expressed ChR2 increases the release probability of neurotransmitter (9) and alters the dynamics of synaptic depression during high frequency stimulation (10), it is necessary to consider a possibility that opsins themselves influence certain forms of synaptic plasticity and compromise the study; however, the information about each specific case remains limited.

One form of short-term synaptic plasticity is the depression of GABA release from GABAergic terminals that undergo repeated stimulation. The GABAb autoreceptors at the terminals mediate that depression, which serves as a feedback control over GABAergic transmission (11–13). In the insular and somatosensory cortex of the rat brain, such depression is particularly pronounced when the interval between stimuli is around 200 ms (14, 15). Recently, we have shown that in brain slices from the mouse dorsomedial prefrontal cortex (dmPFC), GABA release from the somatostatin-positive GABAergic neurons (SOM-INs) expressing ChR2 undergoes strong presynaptic GABAb receptor-mediated depression during their stimulation with the 5Hz frequency train of blue light pulses (6). Here, we compared the GABA depression between two groups of slices prepared from the brains transduced with two different amounts of AAV expressing ChR2, and found that the higher viral quantities diminished the GABAb receptor-mediated depression of GABA release from SOM-IN of dmPFC.

## Materials and methods

### Animals and surgeries

C57BL/6N males were crossed with homozygous 129SvEv somatostatin interneurons-specific Cre-driver females *Sst^tm2.1(cre)Zjh^* (16) to obtain heterozygous *Sst^tm2.1(cre)Zjh^* mice. ChR2-AAV pseudo-type 1 virus containing Cre-activated ChR2 gene was prepared by the University of North Carolina Gene Therapy Vector Core (Chapel Hill, NC) using a plasmid pAAV-EF1a-double floxed-hChR2 (H134R)-EYFP (Addgene #20298). The male progeny at p21-p30 were injected bilaterally into dmPFC at 1.3 mm anterior, 0.4 mm lateral from the bregma, and 1.3 mm ventral from brain surface with 0.5 μl of the viral solution containing either 1 × 10^8^ or 5 × 10^8^ viral particles per hemisphere, as described (5). All experiments were approved by Virginia Tech IACUC and followed the NIH Guide for the Care and Use of Laboratory Animals.

### Electrophysiology

Slice preparation and recordings were performed as described in (6). Coronal dmPFC slices, 300 μm thickness, were prepared from p60-75 mice using DSK Microslicer (Ted Pella, Redding, CA) and ice cold cutting solution contained (in mM) 110 Choline Cl, 2.5 KCl, 1.2 NaH_2_PO_4_, 2.5 NaHCO_3_, 20 glucose, 0.5 CaCl_2_, and 5 MgSO_4_, bubbled with a 95% O_2_/5% CO_2_. Before recording, slices were incubated at least 1 h at room temperature in (in mM) 120 NaCl, 3.3 KCl, 1.0 NaH_2_PO_4_, 25 NaHCO_3_, 10 glucose, 0.5 CaCl_2_, and 5 MgSO_4_, bubbled with 95% O_2_/5% CO_2_. Recording chamber was superfused at 2 ml/min with (mM) 120 NaCl, 3.3 KCl, 1.0 NaH_2_PO_4_, 25 NaHCO_3_, 10 glucose, 2 CaCl_2_, and 1 MgCl_2_, equilibrated with 95% O_2_/5% CO_2_. Whole cell recordings were obtained at 30±1 °C with Multiclamp 700B amplifier and Digidata 1440A (Molecular Device, Sunnyvale, CA). Recordings were performed from the dmPFC region located within +1.9 to +1.3 mm from bregma and 0.5 to 1.5 mm below the brain surface, which includes the prelimbic and anterior cingulate areas. Putative layer V principal neurons were identified by their pyramidal morphology under the Dodt gradient contrast optics (custom made) at 850 nm LED illumination (Thorlabs, Newton, NJ) and were recorded using 4-6 MΩ pipettes filled with (in mM): 120 Cs-methanesulfonate, 5 NaCl, 1 MgCl_2_, 10 HEPES, 0.2 EGTA, 2 ATP-Mg, 0.1 GTP-Na, 5 QX314, pH 7.3, osmolarity 285 Osm. Series resistance (Rs) was 10–20 MΩ and monitored throughout experiments. Data were not included in the analysis if Rs changed more than 20%. All membrane potentials were corrected for the junction potential of 12 mV. Light pulses (470 nm, 1 ms) were generated using an LED lamp (Thorlabs) and a custom LED driver based on MOSFET and were delivered through a 40x objective lens (Olympus, Center Valley, PA) at 0.3–2.5 mW, calibrated by a photodiode power sensor (Thorlabs) at the tip of the lens. In most experiments, data from each cell were obtained from 19 to 20 stimuli sweeps separated by 20 s intervals. CGP52432 was from Tocris (Bristol, UK) and remaining chemicals were from Sigma-Aldrich (St. Louis, MO).

### Data analysis

Statistical analyses were performed using GraphPad Prism (GraphPad Software, La Jolla, CA) and StatView (SAS Institute, Cary, NC). Differences were tested using the two-tailed paired t-test and repeated measure ANOVA, and deemed significant with p<0.05.

## Results

The short-term plasticity of GABAergic synapses formed by the somatostatin interneurons (SOM-INs) on the layer V principal neurons (PN) was examined by recording inhibitory postsynaptic currents (IPSCs) evoked in the PNs voltage-clamped at 0 mV. The IPSCs were evoked by stimulating SOM-INs expressing ChR2 with 5 pulses of blue light at 5 Hz frequency. (Fig.1A, C). The light intensity was adjusted to obtain maximum IPSC responses. A comparison was made between slices expressing “high” and “low” levels of ChR2-YFP, which were obtained from animals transduced with 1 × 10^8^ (“low virus”) and 5 × 10^8^ (“high virus”) viral particles per hemisphere. The examples of ChR2-YFP fluorescence in dmPFC of animals transduced with the two levels of virus are shown on Fig. 1B.

**Figure 1.**
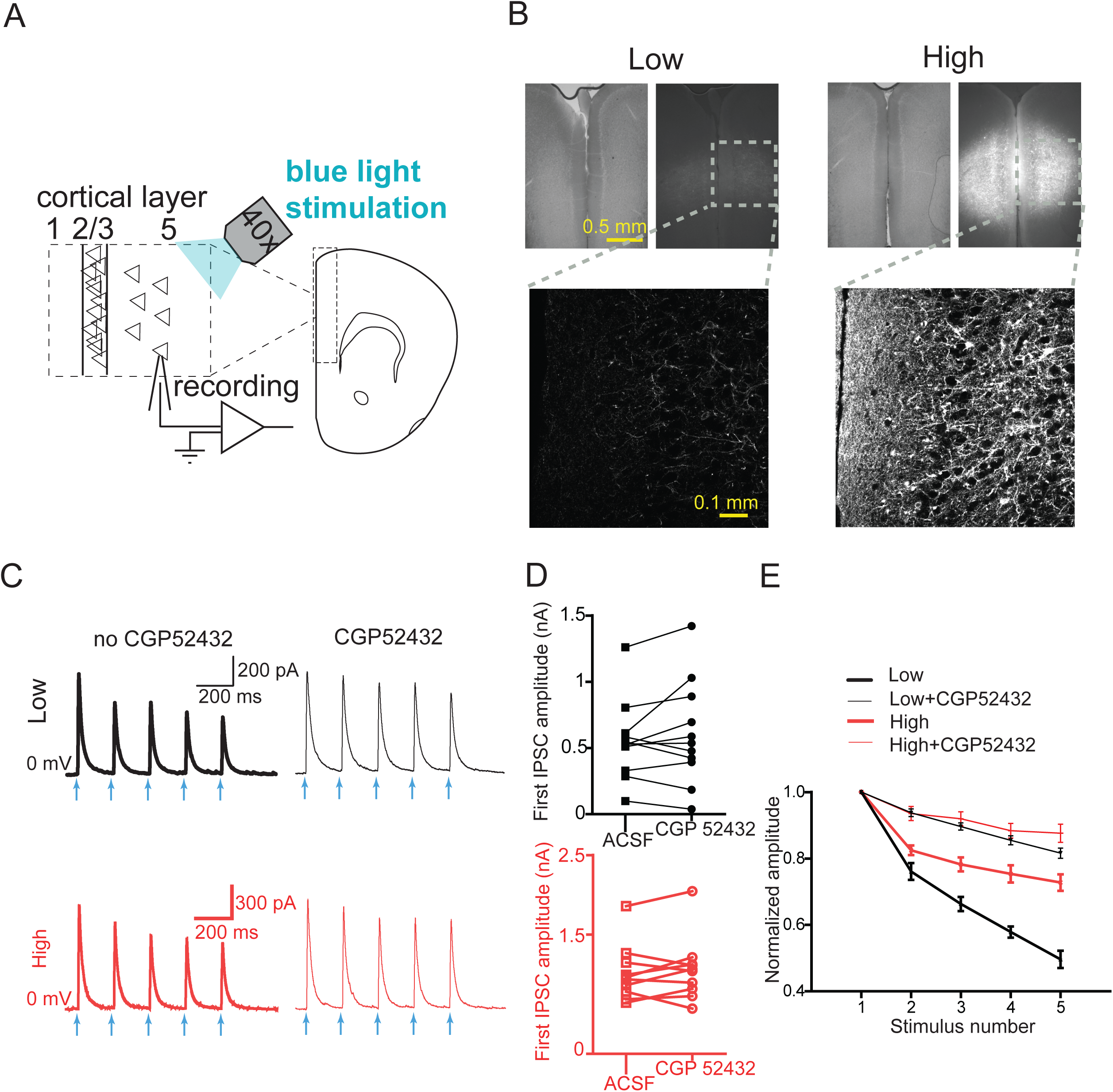
ChR2 attenuates GABAbR-mediated depression of GABA release. A) Experimental scheme. Blue light stimulation of dmPFC and recording from layer V PCs are illustrated. B) dmPFC slices expressing ChR2-YFP in SOM-interneurons transduced with the low (Low) and high (High) amounts of AAV with ChR2-YFP gene. Upper: visible (left) and yellow fluorescence (right) images of the same slices. Lower: confocal images of the areas marked with dashed squares. C) Examples of IPSCs evoked in layer V PCs by 5 Hz train of blue light stimulation of SOM-interneurons expressing ChR2 before and after perfusion with CGP52432 (10 μM), averages of 5 sweeps are shown. The light pulses are indicated by blue arrows. D) Amplitudes of the first IPSCs in the train before and after perfusion with CGP52432, lines connect data points representing the same cells. E) IPSCs amplitudes normalized to the values of the first IPSC, before and after perfusion with CGP52432. Black and red colors represent slices with low (n=11 cells, 3 mice) and high (n=10 cells, 3 mice) levels of viral transduction.

To examine the role of GABAb receptor in the synaptic plasticity, IPSCs were recorded first in ACSF without drugs and then, after 10 min perfusion with 10 μM of CGP52432, a GABAb receptor blocker. The IPSC data from the low virus slices are a part of an earlier publication (6), but are included here to allow for the comparison between the high and low virus conditions.

The IPSC amplitudes decreased along the train in both the low and the high virus groups (low virus: F(4,10) = 121, p<0.0001; high virus: F(4, 9) = 82, p<0.0001), but the decreases in the low virus group were greater (“train” * “level of virus” interaction, F (4,19) = 17.4, p<0.0001) (Fig.1E). CGP52432 did not alter the amplitudes of IPSCs evoked by the first pulse in the train (Fig.1D), but attenuated the IPSC depression in both groups (“train” * CGP52432 interaction, low virus: F(4,20) = 39.4, p<0.0001; high virus: F(4, 18) = 11.0, p<0.0001). The attenuation was greater in the low virus group (“train” * CGP52432 * “level of virus” interaction: F(1,4,38) = 6.23, p=0.0001). Furthermore, CGP52432 eliminated the differences in IPSC attenuation along the train between the low and high virus groups (F(4,19)=2.48, p>0.05). Together, the data indicate that the GABAb receptor signaling is mainly responsible for the differences in IPSC depression between the high and low virus groups.

## Discussion

The key finding here is that with the higher expression of ChR2 in SOM-INs, the light-induced GABA release from these cells is less susceptible to the GABAbR-mediated depression. The GABAbR signaling suppresses neurotransmitter release by inhibiting the adenylate cyclase and voltage gated calcium channels at presynaptic terminals (11). Given that ChR2 conducts Ca2+ (17), it is likely that high levels of ChR2 provide a sufficient Ca2+ influx to override the effect from inhibition of the voltage-gated calcium channels. Another possibility is that excessive levels of ChR2 in the plasma membrane could interfere with normal assembly of the membrane-bound components of the second messenger signaling pathways and thereby make presynaptic terminals refractory to certain forms of neuromodulation.

However, our findings do not imply that overexpression of ChR2 always overrides any presynaptic inhibitory mechanisms of neurotransmitter release. For example, activation of the dopamine receptor 2 in PV-INs and SOM-INs in the basolateral amygdala transduced with 5 × 10^8^ viral particles of the identical virus per hemisphere, which corresponds to the high amounts of virus in the present study, significantly attenuated blue light-induced GABA release from both types of interneurons (3). The main implication of our observation is that using ChR2 as a probe for synaptic modulation carries a risk of interfering with the modulation per se; however, such artifacts can be minimized by decreasing the amounts of ChR2 expression.

## Acknowledgements/Funding Sources

The study was supported by the NIH grant MH112093.

